# Effect of caffeine and other xanthines on liver sinusoidal endothelial cell ultrastructure

**DOI:** 10.1101/2023.01.20.524909

**Authors:** Hong Mao, Karolina Szafranska, Larissa Kruse, Christopher Holte, Deanna L. Wolfson, Balpreet Singh Ahluwalia, Cynthia B. Whitchurch, Louise Cole, Glen P. Lockwood, Robin Diekmann, David Le Couteur, Victoria C. Cogger, Peter A.G. McCourt

## Abstract

Xanthines such as caffeine and theobromine are among the most consumed psychoactive stimulants in the world, either as natural components of coffee, tea and chocolate, or as food additives. The present study assessed if xanthines affect liver sinusoidal endothelial cells (LSEC). Cultured primary rat LSEC were challenged with xanthines at concentrations typically obtained from normal consumption of xanthine-containing beverages, food or medicines; and at higher concentrations below the *in vitro* toxic limit. The fenestrated morphology of LSEC were examined with scanning electron and structured illumination microscopy. All xanthine challenges had no toxic effects on LSEC ultrastructure as judged by LSEC fenestration morphology, or function as determined by endocytosis studies. All xanthines in high concentrations (150 μg/mL) increased fenestration frequency but at physiologically relevant concentrations, only theobromine (8 μg/mL) showed an effect. LSEC porosity was influenced only by high caffeine doses which also shifted the fenestration distribution towards smaller pores. Moreover, a dose-dependent increase in fenestration number was observed after caffeine treatment. If these compounds induce similar changes *in vivo*, age-related reduction of LSEC porosity can be reversed by oral treatment with theobromine or with other xanthines using targeted delivery.

## 1. Introduction

Coffee is one of the most widely consumed beverages meaning that any potential effects related to coffee intake will have significant global implications. Furthermore, most people drink coffee daily due to its stimulating effects. The main active ingredient in coffee – caffeine, can be found also in other beverages (tea, soft and energy drinks), food (guarana berries, chocolate), dietary supplements or even painkillers [1]. Formulations consisting of caffeine and either aspirin (a non-steroidal anti-inflammatory drug, NSAID) or paracetamol, are effective treatments for headaches [2] and sore throats [3] and caffeine has been shown to have an analgesic effect alone [4]. Many studies have investigated the associations between caffeine or coffee intake and health/diseases. Recently, the correlation between increased coffee consumption and improved neurocognitive functioning parameters was elicited in patients infected with human immunodeficiency virus (HIV) and hepatitis C virus (HCV) [5]. A different study showed that a sub-group of patients suffering from multiple sclerosis (MS) benefitted from additional coffee intake [6]. Interestingly, in a meta-analysis of studies about the effects of caffeine on Parkinson’s disease (PD) caffeine appeared to both lower the rate of PD progression in patients suffering from the disease as well as decrease the risk of developing PD in the healthy population [7]. On the other hand, several studies have noted that an adjustment may be needed for an individual’s caffeine dosage in relation to their age, sex and health conditions to maximize the positive and limit negative effects [8][9].

Caffeine is primarily metabolized in the liver to theobromine, theophylline and paraxanthine (Figure 1). The demethylation process occurs in the hepatocytes via the cytochrome P450 enzyme system [10]. Overall, more than 25 metabolites have been identified in humans deriving from caffeine metabolism [11][12], with paraxanthine accounting for approximately 80% of the metabolites in the human liver [13][14]. In rats and mice, however, all three main metabolites of caffeine are present in similar amounts [15][16]. All the aforementioned primary metabolites are pharmacologically active and their plasma concentration may exceed that of caffeine in some stages due to their respective rate of metabolism and clearance [10], especially paraxanthine [16][17].

**Figure 1.**
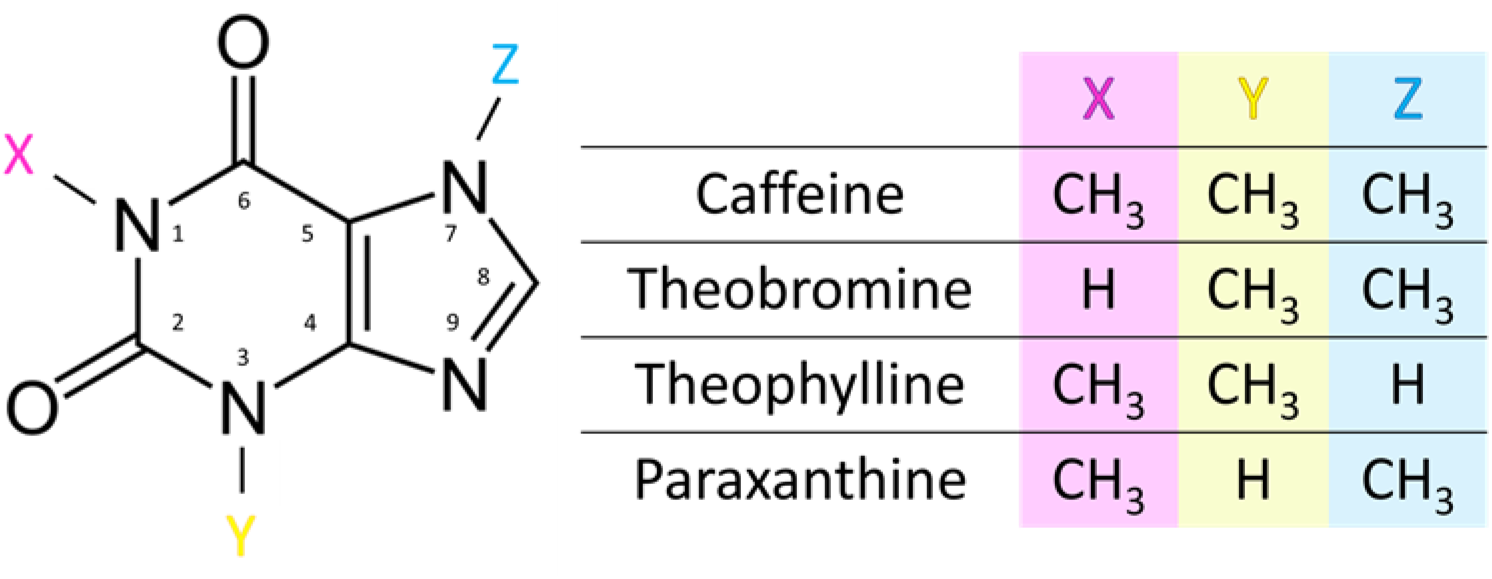
Chemical structure of xanthines. Caffeine (1,3,7-trimethylxanthine) is metabolized by demethylation in the liver to 3 main compounds: theobromine (3,7-dimethylxanthine), theophylline (1,3-dimethylxanthine) and paraxanthine (1,7- dimethylxanthine) [10].

The liver, the body’s largest internal organ, has the major role of detoxifying various metabolites in the human body. Liver sinusoidal endothelial cells (LSEC), which are located in hepatic sinusoids, play a fundamental role in maintaining the homeostasis and metabolic integrity of the liver [18]. LSEC are the most effective scavengers of blood-borne waste [19], regulate sinusoidal blood flow [20], trigger liver regeneration and contribute to hepatic complications, such as liver fibrosis and liver metastasis [21][22][18]. The distinct dynamic morphological features of LSEC – called fenestrations/fenestrae, enable bidirectional size-based transfer between the blood and the underlying hepatocytes. Fenestrations are non-diaphragmed nano-pores with diameters of 50-350 nm [23][24], and they are typically grouped in clusters called sieve plates [25][26]. The structural integrity of fenestrations is vital for the maintenance of regular exchange between the liver and the blood, and alterations in fenestrations can affect hepatocytes and liver function [26]. The regulation of fenestration size and frequency is not yet fully understood, but some signaling pathways have been shown to influence the formation of fenestrations [24]. For example, sildenafil (the active constituent of Viagra) was shown to improve porosity in both young and old mice [27], presumably via regulation of intracellular cGMP levels (via nitric oxide) and/or vascular endothelial growth factor (VEGF)-related/(VEGF)-independent pathways [28][29]. It should be noted however that other pathways may also be involved in the regulation of fenestration size and number (e.g., those mediated by calcium, regulated by myosin light chain (MLC) [24] or by alpha/beta-adrenergic receptors among others [30]).

Even though caffeine is primarily metabolized in hepatocytes, LSEC regulate transportation between the plasma and hepatocytes. Fenestrations create a dynamic barrier that can adapt to environmental conditions in a matter of seconds [31]. Caffeine intake is generally considered to be safe in moderate amounts (≤ 400 mg /day) in healthy adults [32][33], and with normal consumption the plasma concentrations are usually between 2-10 μg/mL (approximately 10-50 μM), rarely exceeding this [34]. Toxic effects occur for plasma concentrations of > 40 μg/mL [34][35]. The adverse effects of large doses of caffeine have been noted from early times, and these include nervousness, anxiety, insomnia, irregular heartbeats, excess stomach acid and heartburn [36]. However, it is important to understand that the liver can be exposed to far greater concentrations of caffeine and related compounds because the uptake takes place in the gastrointestinal (GI) tract. All the venous blood from the GI tract is collected into the portal vein, which then provides 75% of the blood inflow to the liver. This is the first-pass effect whereby the initial local concentration in the liver can be much higher while systemic plasma concentrations of the studied compound are lower [37].

Commercially available products rarely lead to toxic plasma concentrations of xanthines. A single standard cup of black coffee can yield a peak plasma caffeine concentration of about 2 μg/mL, while caffeine supplements can yield up to 10 μg/mL [38]. However, somewhat higher concentrations (8-20 μg/mL) are needed for the therapeutic effect of the caffeine metabolite theophylline in controlling asthma in adult humans [39][40]. Life-threatening effects were reported for the theophylline serum concentrations above 30 ug/mL [41]. Chocolate contains two methylated xanthine derivatives (caffeine and theobromine), which may contribute to its reinforcing effects [42]. It was reported that 370 mg theobromine from chocolate (40-80g of dark chocolate or 110g of regular milk chocolate, roughly the size of a chocolate bar) was rapidly absorbed in humans and produces a plasma concentration of 8 μg/mL after 2 hours [43][44][45]. In humans, paraxanthine plasma levels are usually higher than caffeine and can remain at a high level for a longer time due to the slower clearance and ongoing metabolism of caffeine [17].

Although coffee, chocolate, and other caffeinated substances such as tea are widely used all over the world, the effects of caffeine and its metabolites on LSEC have not been investigated. Here, we study the influence of xanthines on rat LSEC in both physiologically achievable concentrations as well as in higher concentrations that can simulate the first-pass effect.

## 2. Materials and Methods

### 2.1. Rat LSEC production and cell culture

Sprague Dawley male rats (Animal Resource Centre, Murdoch, Western Australia; Charles River Laboratories, Sulzfeld, Germany) were kept under standard conditions and fed standard chow ad libitum (Glen Forrest, Western Australia; RM1-E, Special Diet Service, UK). The experimental protocols were approved by the ethics committee of the Sydney Local Health District Animal Welfare Committee (Approval 2017/012A) and National Animal Research Authority at the Norwegian Food Safety Authority (Mattilsynet; Approval IDs: 4001, 8455, and 0817). All experiments were performed in accordance with relevant approved guidelines and regulations.

Rats (body weight 300-400 g, age 2-3 months) were anesthetized with a mixture of 10 mg/kg Xylazine (Bayer Health Care, California, USA) and 100 mg/kg ketamine (Ketalar, Pfizer, New York, USA), and LSEC were isolated and purified as described Smedsrød et al. [46]. Several fractions of cells were frozen and prepared as described previously [47].

Reagents included caffeine (Cat No. C0750; Sigma-Aldrich, Oslo, Norway), theobromine (Cat No. T4500; Sigma-Aldrich, Oslo, Norway), theophylline (Cat No. T1633; Sigma-Aldrich, Oslo, Norway), and paraxanthine (Cat No. P6148; Sigma-Aldrich, Oslo, Norway). Fibronectin was isolated from human plasma using Gelatin-Sepharose 4B (Cat No. 17-0956-01, GE Healthcare, Sydney, Australia) according to the manufacturer’s instructions. All reagents were dissolved in serum-free RPMI media (Sigma-Aldrich, Sydney, Australia; Oslo, Norway). All experiments were performed in triplicate, using cells isolated from 3 different rats. *In vitro* treatment of LSEC with caffeine was at 8 and 150 μg/mL. Metabolites of caffeine: theobromine (8 and 150 μg/mL), theophylline [40] (20 and 150 μg/mL), paraxanthine (8 and 150 μg/mL) were applied at physiologically relevant and high concentrations in the same manner as for caffeine. For the dose-response study, rat LSEC were treated with caffeine at concentrations: 1, 8, 50, 150, 250 and 500 μg/mL.

### 2.2. Scanning electron microscopy (SEM)

For SEM preparation, the thawing and LSEC culturing protocols are described elsewhere [47]. Cells were plated on 0.2 mg/mL fibronectin coated coverslips and cultured (37°C, 5% CO_2_) for 3 h in serum-free RPMI-1640 (with 10,000 U/mL Penicillin, 10 mg/mL Streptomycin, 1:100) (Sigma-Aldrich, Sydney, Australia) at a density of 0.2 × 10^6^ cells/cm^2^. The LSEC were treated with various agents for 30 min to determine their effects on fenestrations, then fixed using McDowell’s solution (4 % formaldehyde and 2.5 % glutaraldehyde in PHEM buffer pH 7.2) for 15 min and stored in McDowell’s solution until preparation for SEM. After washes with PHEM, the coverslips containing the cells were treated with freshly made 1% tannic acid in 0.15 mol/l PHEM buffer for 1 hour, 1% OsO_4_ in water for 1 hour, dehydrated in ethanol (30%, 60%, 90% for 5 minutes each, 5 times 100% ethanol for 4 minutes each), and incubated twice in hexamethyldisilazane for 2 minutes (Sigma-Aldrich, Oslo, Norway), before coating with 10-nm gold/palladium alloys. Imaging and image analysis was performed blind to the sample ID. Coded specimens were examined using a commercial SEM (Sigma HV0307 or Gemini 300, Zeiss) at 2kV. Large fields of view (magnification of 1k) containing several/individual cells were randomly acquired to determine cell culture condition and cell size, and high-resolution SEM images of cell at magnification of 15k were selected blindly to assess fenestration size.

Fenestration size, porosity and frequency were assessed in SEM images from each cell culture selected from different areas. Open pores with diameters between 50–350 nm were defined as fenestrations and holes larger than 350 nm as gaps. Porosity was defined as the sum area of fenestrations per total area of the cell in the micrographs. Frequency was denoted as the total number of fenestrations per sum area of the cell excluding the sum area of gaps [48].

### 2.3. Structured illumination microscopy (SIM)

After fixation, the cells were stained with CellMask Green (1:1000 in phosphate buffered saline (PBS)) for 10 minutes, washed 3 times in PBS and then mounted in Vectashield antifade mounting medium (Vectro Laboratories, Burlingame, California) and imaged using a commercial super-resolving SIM (DeltaVision/OMXv4.0 BLAZE, GE Healthcare) with a 60X 1.42NA oil-immersion objective (Olympus). 3D-SIM image stacks of 1 μm were acquired with a z-distance of 125 nm and with 15 raw images per plane (five phases, three angles). Raw datasets were computationally reconstructed using SoftWoRx software (GE Healthcare). The datasets were further analyzed using the pixel classification workflow in the freely available machine learning image processing software Ilastik [49]. Fenestration detection steps were described in our previous study [50]; notably, the detected objects with diameters below 50 nm and above 300 nm were excluded prior to binning for SIM images.

### 2.4. Endocytosis and degradation assay

For quantitative studies of endocytosis and degradation, fully confluent cultures of rat LSEC (approx. 0.25–0.3 × 10^6^ cells/cm^2^) were established in 48-well culture dishes coated with fibronectin, were pretreated with different agents for 30 minutes (at 37 °C, 5% CO_2_), subsequently incubated in 0.2 mL serum-free RPMI containing 1% human serum albumin and ∼20000 cpm ^125^I-FSA for 2 h (total incubation time with drugs was 2.5 h). Thereafter, the cell-associated and degraded FSA fractions were analyzed as described previously [51][52]. Briefly, LSEC scavenge ^125^I-FSA via the endocytic cell surface receptors stabilin-1 and -2, and this is later digested, resulting in the separation of FSA and ^125^I (free iodine). The degraded fraction was calculated from the measurement of free iodine in the supernatant solution after precipitation of the remaining intact ^125^I-FSA using 20% trichloroacetic acid. The cell-associated fraction was measured after lysis of the cells with 1% sodium dodecyl sulfate (SDS). The data was analyzed after subtraction of the unspecific binding and free iodine content in the cell free controls. The radioactivity was measured using a Cobra II, Auto-Gamma detector (Packard Instruments, Laborel, Oslo, Norway).

### 2.5. Data analysis and statistics

We selected image analysis methods according to our previous study [48]. Fenestration diameters were measured from SEM images using a threshold-based semi-automated method. This approach allowed us to greatly improve both the number of analysed fenestrations as well as the precision of measurement. Mean fenestration diameters were calculated from the single fenestration areas obtained from the segmented images (See supplementary information Figure S2). Fenestration frequency was obtained from the manually counted fenestrations from each cell. This reduced the influence of any imaging artifacts including differences in image quality between the samples. Porosity was calculated as a combination of both fenestration number and size distribution (details can be found in the supplementary information of [48]).

All graphs were prepared using OriginPro software (OriginPro 2021, OriginLab Corp., Northampton, Massachusetts) and image analysis was performed using the free, open-source software Fiji/Image J [53]. Significance was assessed using a two-tailed student T-Test for porosity and fenestration frequency parameters. Fenestration diameters and endocytosis experimental data were analyzed with one-way ANOVA performed using the GraphPad Prism 8 (La Jolla, USA, www.graphpad.com), and the post hoc tests were used with Dunnett’s multiple comparisons test for each treatment with control. The results were considered significant if p<0.05 (*) or described as trends for p<0.1 (#).

## 3. Results and discussion

### 3.1. Dose-response of LSEC treated with caffeine

A typical flat, well-spread morphology was observed in all samples, and cell culture purity was assessed based on the presence of the unique morphological feature of LSEC - fenestrations (Figure 2). SEM data with the analyzed number of cells as well as fenestration diameter, porosity (i.e. percentage of the cell surface covered by fenestrations) and frequency (i.e. the number of fenestrations per cell area) following various treatments are summarized in Table 1. The fenestration diameter after treatment remains within the range of 160 to 180 nm, which is consistent with previously reported data [23][24]. Roughly 20000- 50000 “holes” were analyzed for each treatment group to calculate fenestration diameters. Fenestration frequency and porosity were measured for ∼35 cells total from 3 individual animals.

**Table 1.** Changes in the parameters describing fenestrated morphology of freshly isolated LSEC after treatment with caffeine. Measurements were extracted from SEM images using semi-automatic (diameter) and manual methods (frequency).

| Concentration<br>[µg/mL] | No. fenestrations<br>measured | Diameter [nm]<br>± HWHM | Porosity<br>[%]<br>± SD | Frequency<br>[No./µm²]<br>± SD | No. cells<br>measured |
| --- | --- | --- | --- | --- | --- |
| Control | 19811 | 176±33 | 3.13±1.23 | 1,25±0.49 | 35 |
| 1 | 35999 | 178±35 | 3.74±2.28 | 1,62±0.99 | 35 |
| 8 | 51571 | 177±36 | 5.03±2.04** | 1,97±0.80** | 33 |
| 50 | 46041 | 170±36* | 4.84±2.68** | 2,09±1.16** | 34 |
| 150 | 52370 | 166±33* | 5.57±1.87** | 2,42±0.81** | 32 |
| 250 | 41741 | 170±34* | 6.77±1.93** | 2,85±0.81** | 34 |
| 500 | 23998 | 179±36 | 6.39±2.10** | 2,54±0.83** | 34 |

SD – standard deviation. \*\* $p < 0.01$ , \* $p < 0.05$

**Figure 2.**
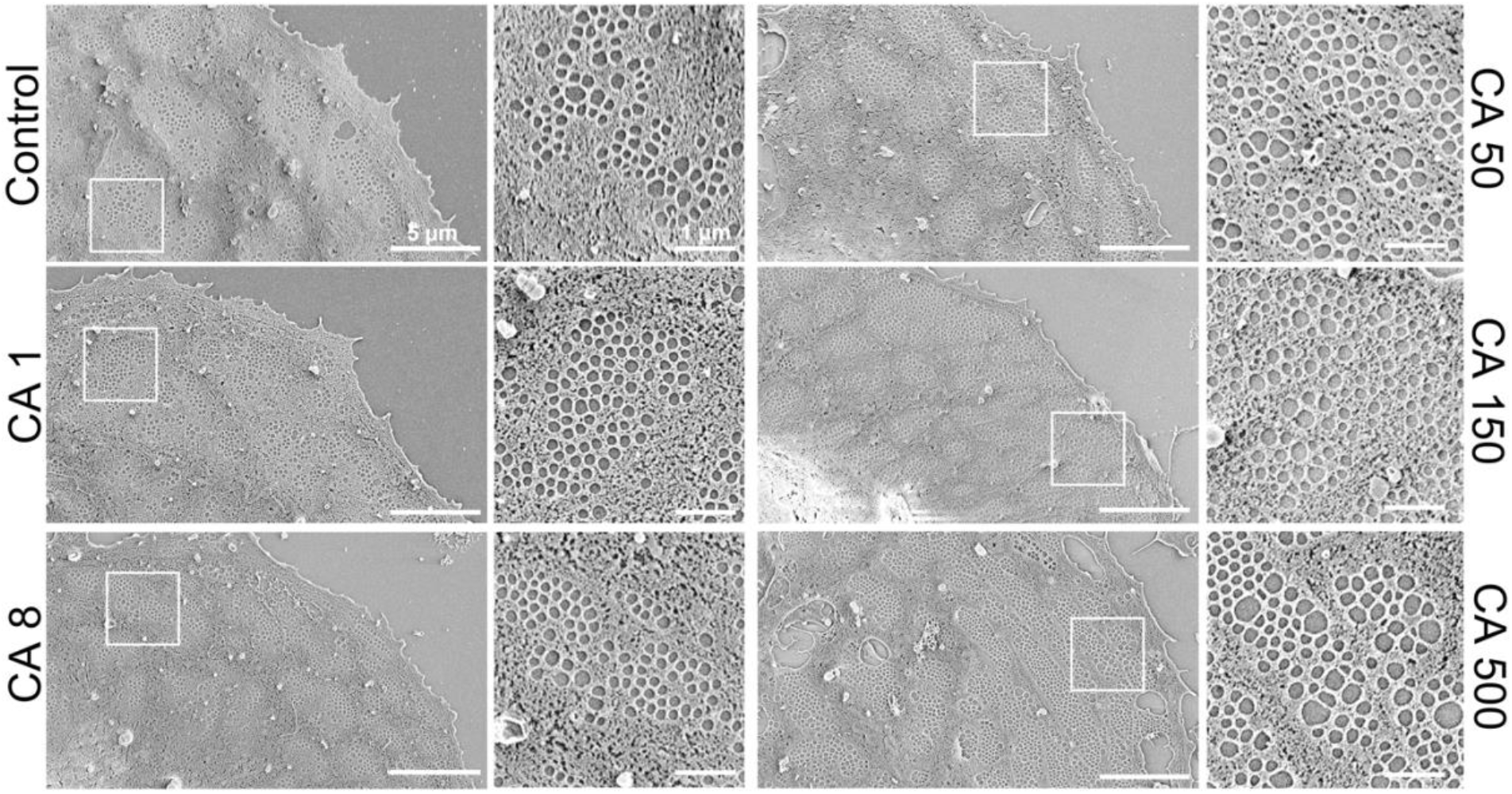
Representative scanning electron microscopy (SEM) images following 30 min of caffeine treatment of rat liver sinusoidal endothelial cell (LSEC) fenestration. Scale bar size 5 μm. CA – caffeine, indicated concentrations are μg/mL.

Changes in fenestration size in combination with differences in fenestration number may also influence the average cell porosity. Fenestration frequency is a parameter describing the number of fenestrations per cell which in combination with fenestration size can be translated into porosity – the percentage of the cell area covered in fenestrations. The diameter of fenestrations is responsible for (size-dependent) selectivity of passive transport between the bloodstream and hepatocytes, while the number of fenestrations per cell/area relates to the rate of liver filtration [54].

To better understand the effect of caffeine, a dose-dependent study was performed using SEM images. A decrease in fenestration diameter was observed for concentrations of 50-250 μg/ml, reaching the minimum at 166 nm at 150 μg/ml (in comparison with 176 nm in the untreated cells) (Table 1, Figure 3A). For the highest used concentration of 500 μg/ml, a small increase in the diameter was observed. The analysis of the fenestration number showed a dose-dependent increase in the fenestration frequency for caffeine concentrations of 8 μg/ml or above (Figure 3B). The highest value of 2.9 fenestrations per μm^2^ was reached at the caffeine concentrations of 250 μg/ml, which is an over two-fold increase compared with the control. No further increase in fenestration frequency was observed. Similar trends were observed for the porosity parameter calculated from the combination of fenestration frequency and diameter data (Figure 3C). Interestingly, the comparison of porosity data for different treatments shows a connection between fenestration size and number. For example, no change in fenestration frequency was observed between the caffeine concentrations of 8 and 50 μg/ml, while a 7 nm decrease in the mean fenestration diameter led to only 0.2 percentage point decrease in the porosity. Similarly, a 13 nm increase in the mean fenestration diameter between the treatments of 150 and 500 μg/ml resulted in a 0.82 percentage point increase in the porosity.

**Figure 3.**
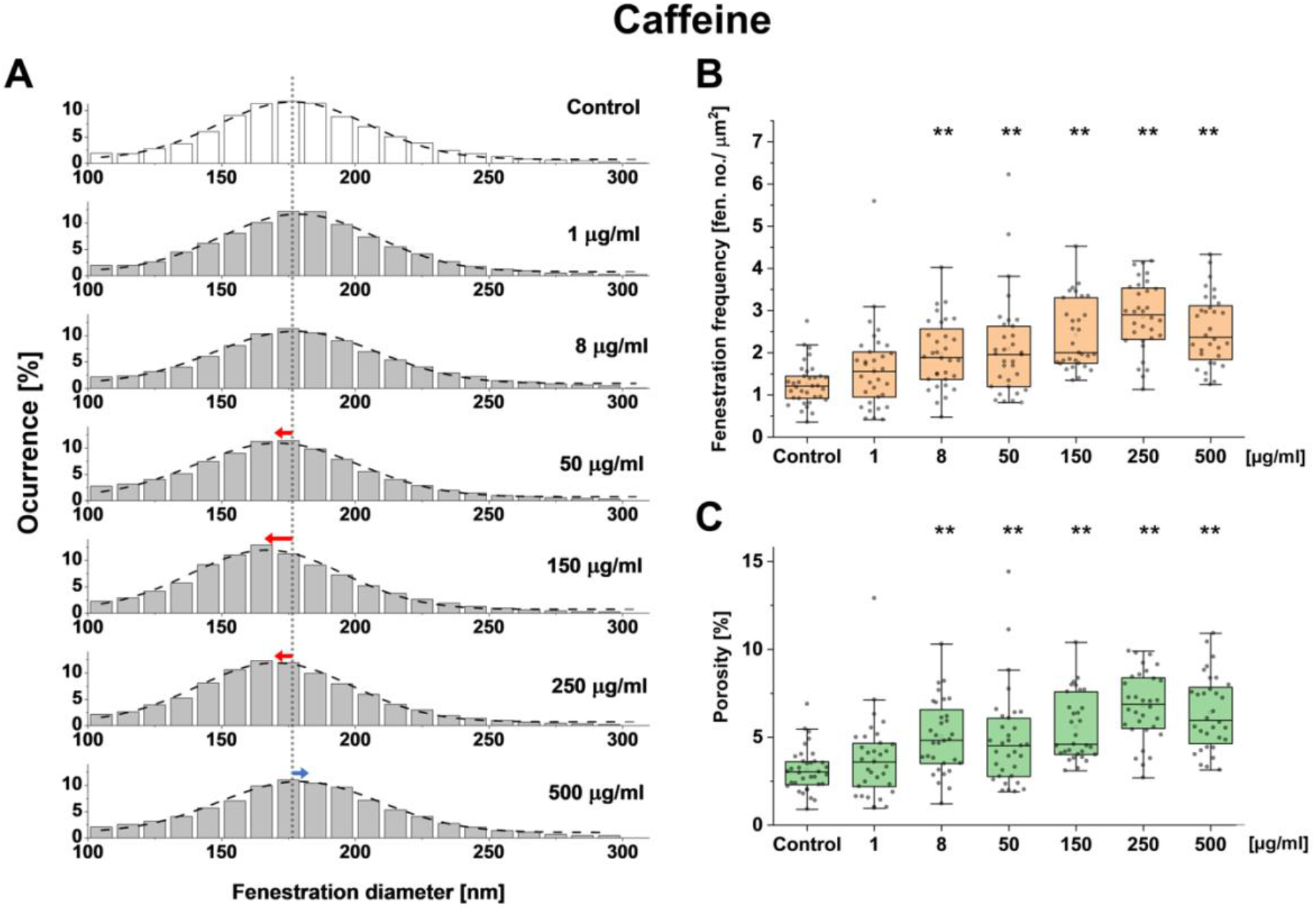
Dose response of LSEC treated with caffeine and changes in fenestration diameter distribution (A), fenestration frequency (B) and porosity (C). (A) dotted line indicates the mean fenestration diameter of the untreated group and arrows show the trends of changes in diameter in the treatment groups. ** p<0.01

### 3.2. The effects of other xanthines on LSEC morphology

Caffeine is metabolized in the liver into three main metabolites: theobromine, theophylline and paraxanthine (Figure 1) [11][12]. Here we study and compare the effects of those xanthines on LSEC. Similarly to the caffeine dose response experiment, we observed typical flat, well-spread morphology in all samples (Figure 4). SEM data with the analyzed number of cells as well as fenestration diameter, porosity and frequency following xanthine treatments are summarized in Table 2. Roughly 7200-69000 fenestrations were analyzed for each treatment group to calculate fenestration diameters. Fenestration frequency and porosity were measured for around 60 cells total from 3 individual animals.

**Figure 4.**
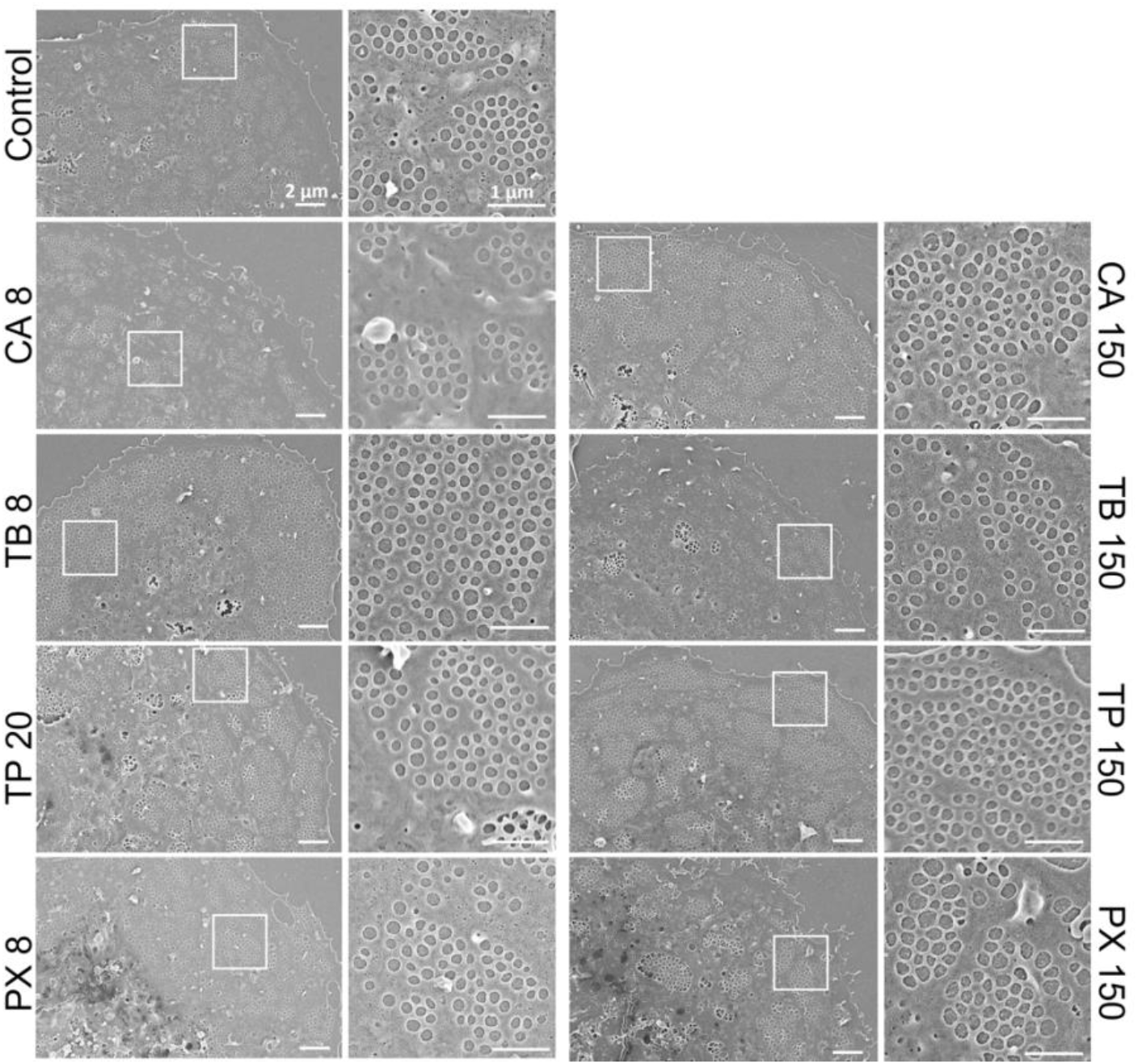
Representative scanning electron microscopy (SEM) images following xanthines treatment of rat liver sinusoidal endothelial cell (LSEC) fenestration. Scale bar size: overview images 2 μm, inset 1 μm. CA - caffeine, TB - theobromine, TP – theophylline, PX - paraxanthine. Indicated concentrations are μg/mL.

**Table 2.** Changes in the parameters describing LSEC fenestrated morphology. Measurements were extracted from SEM images using semi-automatic (diameter) and manual methods (frequency).

| Treatment |  | No. fenestrations measured | No. cells | Diameter [nm] ± SD | Porosity [%] ± SD | Frequency [No./µm<sup>2</sup>] ± SD | No. cells measured |
| --- | --- | --- | --- | --- | --- | --- | --- |
| Drug | Concentration [µg/mL] |  |  |  |  |  |  |
| Control | - | 68829 | 157 | 190 ± 51 | 7.50 ± 4.24 | 2.52 ± 1.33 | 51 |
| Caffeine | 8 | 7235 | 33 | 176 ± 46* | 7.04 ± 3.27 | 2.72 ± 1.27 | 49 |
|  | 150 | 18844 | 34 | 179 ± 53* | 9.13 ± 3.91* | 3.07 ± 1.38* | 60 |
| Theobromine | 8 | 12771 | 20 | 185 ± 48* | 7.50 ± 3.29 | 3.32 ± 1.42* | 55 |
|  | 150 | 7029 | 17 | 177 ± 58* | 7.65 ± 3.48 | 3.15 ± 1.43<sup>#</sup> | 49 |
| Theophylline | 20 | 13496 | 27 | 188 ± 51* | 8.15 ± 3.36 | 2.61 ± 1.14 | 54 |
|  | 150 | 20311 | 33 | 178 ± 47* | 8.97 ± 3.45<sup>#</sup> | 3.23 ± 1.39* | 60 |
| Paraxanthine | 8 | 15210 | 21 | 178 ± 46* | 8.61 ± 3.71<sup>#</sup> | 2.80 ± 1.28 | 60 |
|  | 150 | 16766 | 36 | 190 ± 55 | 8.79 ± 4.27<sup>#</sup> | 2.88 ± 1.40<sup>#</sup> | 56 |

SD – standard deviation. \*p<0.05, #p<0.1.

The cell morphology from different treatment groups was observed after 30 minutes incubation. No drastic changes and signs of toxicity were noticed. Only the high dose (150 μg/mL) of theobromine showed possible early signs of toxicity in some cells – namely irregular cell edges and shrinking.

### 3.3. Quantitative effects on fenestrations

Some of the xanthines elicited changes in the fenestration frequency (Figure 5, Table 2): a significant increase was observed for the high doses (150 μg/mL) of caffeine and theophylline and lower dose (8 μg/mL) of theobromine, (22%, 28% and 32%, respectively). Also, high doses of theobromine and paraxanthine show increasing trends in fenestration number. The less prominent effect of high versus low dose of theobromine may be a result of early toxic effects suggested by the visual examination of the cell images - some cells treated with the high dose showed signs of irregular cell edges or reduced cell area. The trend after paraxanthine treatment may suggest that exposure to the drug longer than the incubation time used here (30 minutes) may further increase LSEC porosity. In humans, paraxanthine is the main metabolite of caffeine and its concentration in the plasma can reach higher values than caffeine itself and remain at this high level for prolonged periods, up to a few hours [17].

**Figure 5.**
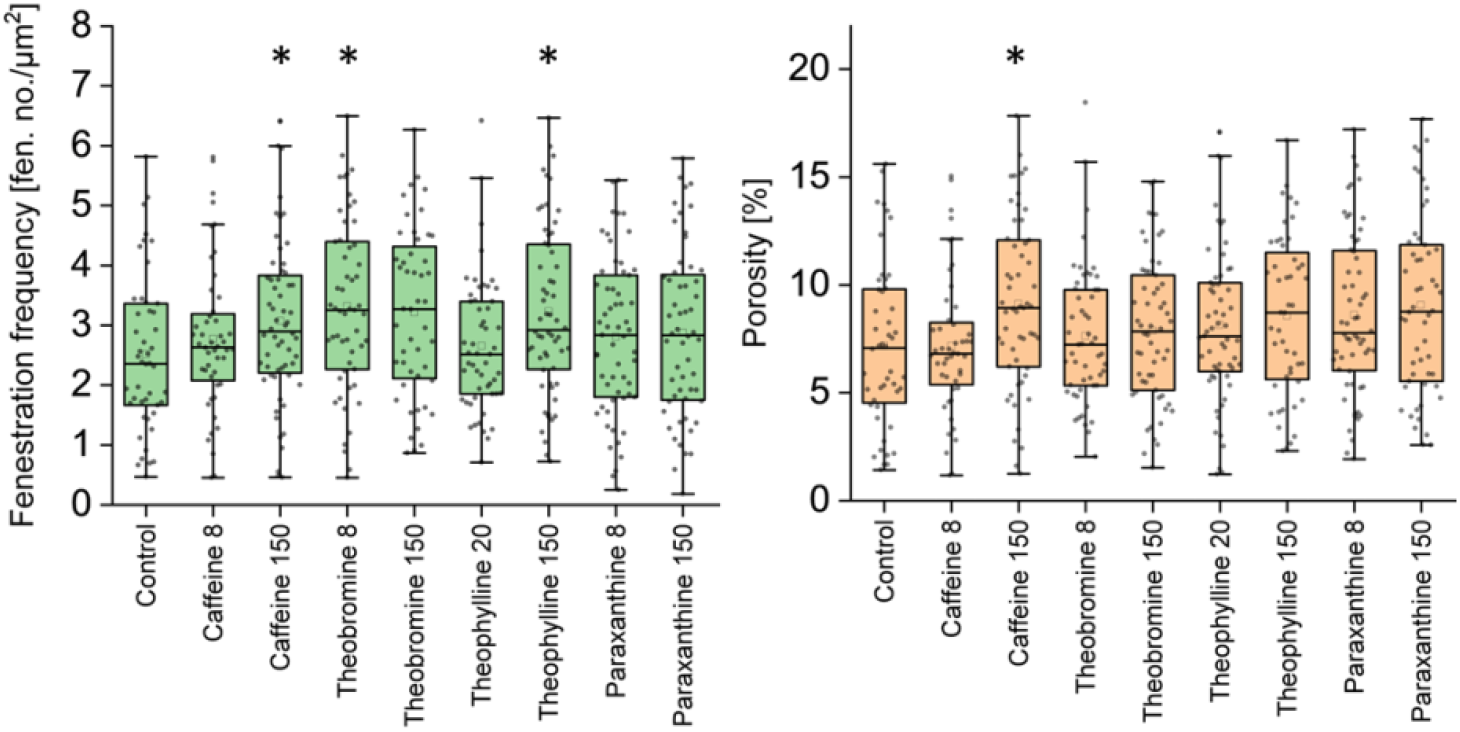
Fenestration frequency and porosity of LSEC treated with xanthines. The graphs present combined data from 3 studied animals. Treatment concentrations are indicated in μg/ml. Each dot represents analyzed SEM image of a single cell. *p<0.05.

Even though we did not observe large changes in the mean fenestration diameters, the porosity parameter shows the influence of the diameter distribution (Table 2). A significant increase (22%) in porosity was observed only in the high dose of caffeine. The high dose of caffeine did not affect the mean fenestration diameter, so the effect is visible in both parameters. On the other hand, high doses of both theobromine and theophylline caused a decrease in fenestration size which counteracted the influence of in-creased fenestration number. *In vivo*, such an occurrence would suggest more rapid filtration of the smaller molecules in the liver. Paraxanthine for both concentrations and high dose theophylline treatments also showed an increasing trend in porosity.

To further understand the changes in fenestration diameter after the various treatments, we studied the whole distribution of fenestration size, instead of just average values. We observed shifts in the diameter histograms, and to better visualize that data we separated fenestrations into 3 size groups: small (S) - <100 nm, medium (M) – 100-200 nm, large (L) - >200 nm. These groups can be related to the filtration of different molecules that are metabolized by hepatocytes – HDL and LDL have a diameter below 100 nm, while chylomicron remnants are usually above 100 nm in diameter [55]. Already in 1970, Wisse pointed out that fenestration size may play a role in liver filtration [20] and later in 1995, Fraser et al. [56] proposed the barrier function of LSEC that is chylomicron remnant clearance, namely that LSEC fenestration diameter dictates which chylomicron remnants can be removed from the circulation. Although the filtration function of LSEC is a combination of both passive filtration and active receptor-mediated trans- and/or endocytosis, the previous studies suggest that transportation of lipoprotein fractions through the endothelial barrier is mostly influenced by the passive flow through the fenestrations [57][58]. Changes in the distribution of fenestrations between those groups relative to control can be found in Figure 6.

**Figure 6.**
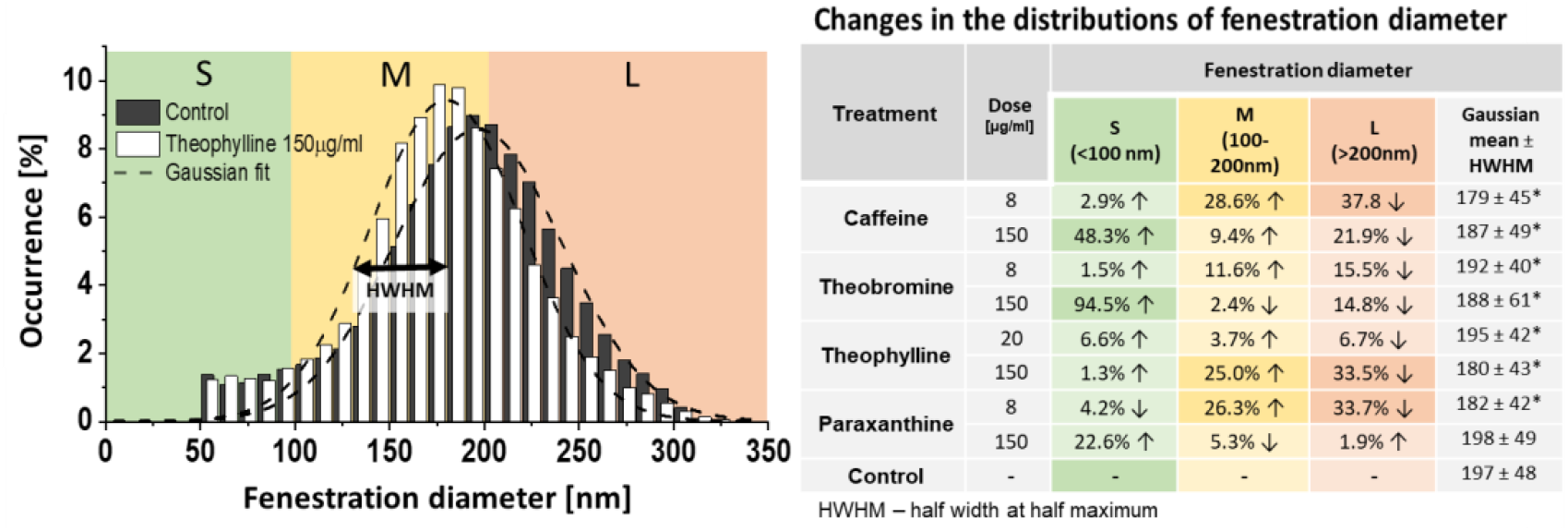
Fenestration distribution in LSEC treated with xanthines. The graph presents the control group and theobromine (high concentration) treatment as an example (all histograms can be found in supplementary materials (Figure S1). S (small)/M (medium)/L (large) represent the fractions of fenestrations according to their size. Changes in the fenestration distribution are presented in the table as relative to the control/untreated samples. Gaussian mean is calculated as a center of Gaussian fit. The data was obtained from SEM images using semi-automated method.

We observed a decrease in the large fenestrations after treatment with all xanthines with the exception of the high paraxanthine dose. The high dose of theophylline and lower dose of paraxanthine showed a 33% decrease in the number of large fenestrations. The high dose of caffeine also was responsible for an almost 22% reduction in the large fenestration number. The medium size fenestration fraction was increased by low doses of caffeine and paraxanthine and a high dose of theophylline by 29%, 26% and 25%, respectively. The remaining treatments show little to no effect in this size range. Interestingly, high doses of caffeine and theobromine cause a large increase in the detectable number of small fenestrations, of 48% and 95%, respectively. The high dose of paraxanthine also resulted in a 23% increase in small fenestrations, but unlike caffeine or theobromine it had very little impact on either medium or large pores.

The detailed analysis of the fenestration size distribution reveals some additional information about the changes in the liver sieve. To date, most studies reported changes in fenestration size as just a difference in the mean value but a Gaussian-like distribution of fenestration was confirmed in multiple studies [26][59][48]. Here, we show that the fenestration distribution shape can remain as a Gaussian distribution with little to no changes to the mean value while changes in specific fractions, reflected in the distribution width, can be significant and have biological implications. For example, low dose paraxanthine results in over 20% increase in <100 nm fenestrations with only 5% increase in medium-size holes without a change in the mean fenestration value – 197 nm and 198 nm in control and after treatment. Meanwhile, such a change *in vivo* can lead to increased availability of VLDL, HDL and other small molecules to hepatocytes, therefore increasing filtration from the bloodstream. On the other hand, a decrease in the large fenestration fraction (such as after low doses of caffeine and paraxanthine or high dose of theophylline) can lead to reduced filtration of some fractions of larger chylomicron remnants. Changes in transportation between plasma and hepatocytes can therefore have both positive and negative effects. Lower filtration can have a protective role against reducing the exposure of some agents to hepatocytes, but it also can in-crease lipoprotein fractions in the blood which may contribute to cardiovascular dis-eases. Wright et al. [60] showed that smaller liver fenestrations observed in rabbits can be a cause of their susceptibility to arteriosclerosis. This theory is supported by the increase of fenestration size in rabbits reported after pantethine treatment, which reduced their sensitivity to dietary cholesterol [61]. The chylomicron remnant size varies from 100 to even 1000 nm, so the reduction in large or medium fenestrations could correspondingly affect/lower the filtration of certain fractions of chylomicrons containing different amounts of triglycerides and cholesterols.

To validate our findings, we also assessed the fenestrations with some treatments on SIM (Figure 7, caffeine 150 μg/mL and theophylline 30 μg/mL are shown for comparison). Similar to the SEM observations, LSECs featured numerous fenestrations which were clustered as sieve plates. Further assessment of the diameter of fenestrations was conducted in a machine learning assisted workflow, as was used in a previous study [50]. By comparing the mean fenestration diameter in our dehydrated SEM result with wet-fixed SIM data, the latter resulted in a 15% smaller size for the control groups [62]. The SIM data confirmed the increase in fenestration frequency and changes in diameter distribution induced by the high dose of caffeine and theophylline (Figure 7).

**Figure 7.**
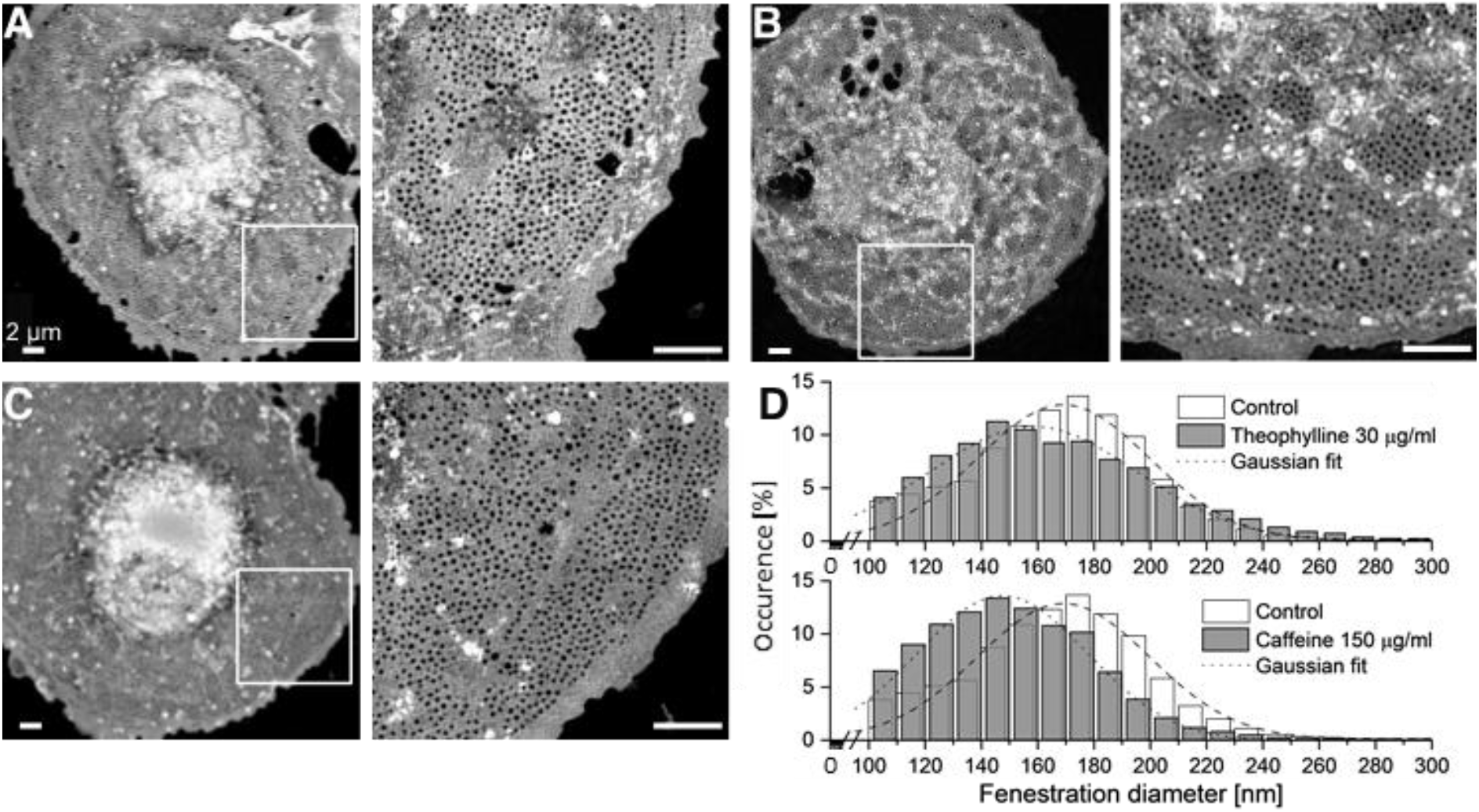
Maximum intensity z-projections of 3D-SIM images show fenestrations grouped in sieve plates (inset) following treatments - A: Control; B: Theophylline (30 μg/mL); C: Caffeine (150 μg/mL); D: Fenestration size distribution after treatments. Mean diameter calculated from Gaussian fits ± HWHM: Control 169±35 nm, theophylline 159±43 nm, caffeine 148±35 nm. Scale bars: 2 μm.

Though not having as high resolution as SEM, the SIM technique gives a resolution double that obtained via conventional light microscopy. The average diameter of fenestrations is well discerned within the regular observed size range. For SIM, the samples can be studied while wet, thus avoiding the artifacts from dehydration required for SEM. 3D-SIM was used in our study to image fenestrations in fixed rat LSECs in Vectashield mounted samples. Importantly, a limitation of the linear SIM is that only fenestrations with a diameter around 100 nm or more are resolved. As mentioned above, by comparing the results of the control group from both methods, the mean fenestration diameter was larger in SEM processed samples, which might be due to the dehydration step during SEM preparation. Notably, due to the resolution limit of SIM being around 100 nm, the diameter analyzed from SIM images was of the same magnitude (Figure 7D).

### 3.4. Xanthines effect on LSEC viability

LSEC are the main scavenger cells in the body, clearing various compounds from the blood [18]. A well-established method for validation of LSEC functional viability is a measurement of their endocytic capacity of denatured or modified proteins such as formaldehyde-treated serum albumin (FSA) [63]. To assess the possible toxic effects of xanthines, we measured the uptake and degradation of radiolabeled FSA in LSEC (Figure 8). The data show no effects on endocytosis after caffeine treatment (up to 1500 μg/ml). Theobromine showed an 8% decrease in total endocytosis at 250 μg/ml, while theophylline and paraxanthine showed 10% and 8% reduction, respectively, at 1500 μg/ml. Evaluation of cell morphology on SEM images did not reveal any apparent morphological disruptions characteristic for LSEC (Figure 4) (indicated by e.g., large gaps >400 nm, cell shrinkage or disrupted cell edges). The additional morphological assessment was performed using SIM (Figure 7). Nevertheless, this small decrease of degradation of FSA could be related to the toxic effects of the compounds. We did not reach *in vitro* toxic concentration of caffeine and the negative effects of other xanthines were observed only at very high concentrations of 250/1500 μg/mL which are above the reported *in vivo* toxic doses [36].

**Figure 8.**
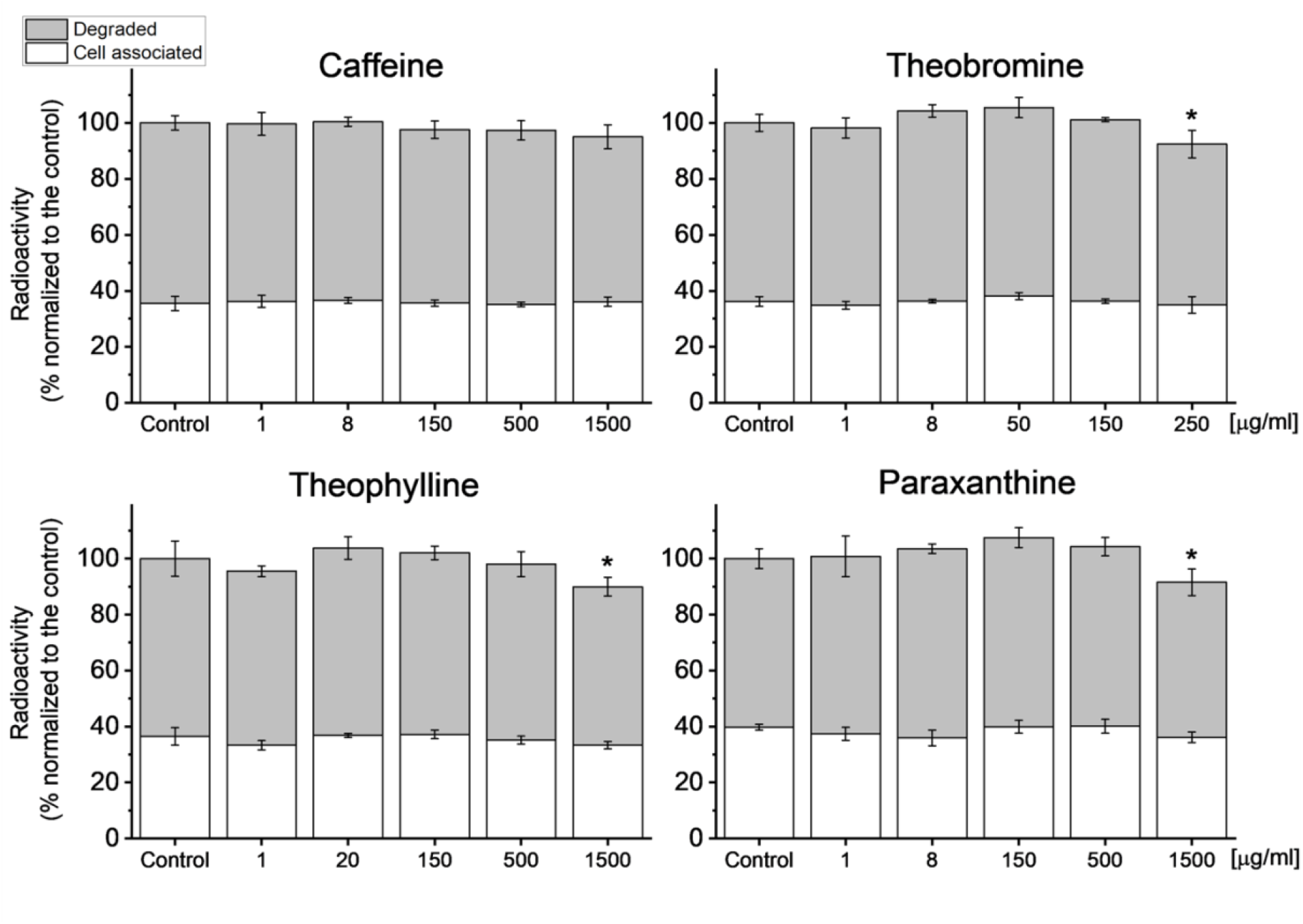
^125^I-FSA uptake in isolated LSEC after treatments with xanthines. Incubation time with ^125^I-FSA – 2h. Bars indicate mean ± SD, n = 3, *p<0.05.

### 3.5. Possible mechanisms of action

Caffeine is metabolized in hepatocytes mainly by the CYP1A2 enzyme and the exons for this enzyme vary between humans [64]. These differences cause variations in caffeine metabolism which affects clearance time and therefore results in the different total stimulating effects of caffeine. LSEC do not express CYP1A2 [65] nor metabolize caffeine per se, however, here we show that xanthines can affect their morphology. The exact mechanism is not known but there are several receptors and pathways present in LSEC that can be affected. Based on our extensive review of mechanisms behind the structure and functioning of LSEC fenestration, some possible mechanisms of action can be proposed [24].

For example, changes in intracellular cGMP can be associated with fenestration regulation. Hunt et al. [27] showed that drugs promoting cGMP and PKG, such as sildenafil, have positive effects on porosity and fenestration frequency. cGMP levels are mainly controlled by the regulation of its degradation by various phosphodiesterases (PDE) as well as by extracellular efflux by ABC transporters [66]. Both caffeine and theophylline were reported to act as non-specific PDE inhibitors [66][67]. These data suggest the possible mechanism of action via the cGMP-PKG pathway, however further studies are necessary to confirm this.

Caffeine and paraxanthine can also act through nonselective antagonizing of adenosine receptors [68], which could influence fenestrations via reduction of cAMP. A decrease of cAMP resulting from serotonin challenge showed a decrease in the fenestration size [30], similar to the observed here after treatment with a high concentration of xanthines. This mechanism has not been confirmed in LSEC yet, however, the effects of adenosine receptors in the liver are currently under investigation [69].

### 3.6. Translational relevance

The physiologically achievable non-toxic plasma concentrations of xanthines in humans are up to about 1-8 μg/mL. The endocytosis data show that xanthines have no negative effects in this concentration range. The decrease in the LSEC endocytic activity was observed only with very high concentrations of theobromine, theophylline and paraxanthine which are not physiologically relevant. The decrease in FSA endocytosis is due to reduced degradation, not uptake, which suggests a possible influence of high concentrations of xanthines on the lysosomal degradation or intercellular transportation, rather than an influence on the scavenging receptors. Together with the LSEC morphology results, it suggests that there is no correlation between the cell porosity and endocytic activity, which agrees with the report of Simon-Santamaria et al. [63].

Daily consumption of 40-80 g of dark chocolate or 110 g of regular milk chocolate yields a plasma theobromine concentration of 4-8 μg/mL in humans, which is within the region that elicits increased porosity in rat LSEC *in vitro*. Our study demonstrated theobromine, at physiologically relevant concentrations, may increase the fenestration frequency in LSEC, which may further affect hepatocyte metabolism and xenobiotic detoxification. If the results in this study translate to the *in vivo* context, the theobromine induced increase in fenestration number could improve liver function by enhancing the bi- directional exchange of substrates between the plasma and the hepatocytes, for example in the elderly who have reduced LSEC porosity. On the other hand, due to fenestration loss associated with ageing, some hepatocyte-targeting drug dosages should be adjusted for elderly people [70]. The otherwise positive effect or re-fenestration (induced reappearance of fenestrations) after xanthine treatment could lead to toxic effects due to the sudden increase of drug levels to which hepatocytes are exposed. This effect may be even more relevant for the people taking multiple drugs at the time as polypharmacy (daily intake of 5 or more drugs) is steadily increasing in recent years [71].

The positive effects of high doses of caffeine and theophylline are however harder to translate into useful intervention for the benefit of liver health as this concentration (150 μg/mL) is within the toxic range for those compounds. Recent studies have suggested the high plasma concentration needed for effect can be avoided if therapies are directly delivered to LSECs via nanoparticles and may result in similar *in vivo* effects as observed *in vitro* [72][73]. Further studies are clearly required to determine if xanthines affect fenestrations *in vivo*. Previous studies investigated the possibility of using rates of caffeine clearance as a guide to deteriorating liver function in cirrhosis. The results could be explained by the effect on LSEC fenestrations as the caffeine clearance did not correlate with conventional liver function tests [37]. These findings suggest that the development of the new tests of hepatic function should take into consideration also possible effects of the agents on the LSEC barrier, and not only hepatocyte metabolism.

## 4. Conclusions

In conclusion, we have shown *in vitro* xanthine treatments with caffeine, theobromine, theophylline and paraxanthine elicit changes in fenestration size, porosity and frequency in rat LSEC. It is only at very high concentrations, that xanthines have an inhibitory impact on the uptake of soluble macromolecules. Caffeine and theophylline in high doses increase fenestration frequency and the number of smaller-sized fenestrations in LSEC. Theobromine at a physiologically relevant dose also increases fenestration frequency in rat LSEC. Although similar findings remain to be confirmed *in vivo*, if theobromine elicits these effects in animal studies, it might prove to be a useful (and simple) intervention (via chocolate) to improve LSEC porosity in elderly people. Concomitant to this, caffeine and theophylline could be used for the improvement of LSEC fenestration number; however, a drug delivery system targeting LSEC would be re-quired to avoid unwanted systemic side effects.

## Author Contributions

Conceptualization, Hong Mao, Karolina Szafranska, Deanna Wolfson and Peter McCourt; Data curation, Hong Mao and Karolina Szafranska; Formal analysis, Hong Mao, Karolina Szafranska, Larissa Kruse and Christopher Holte; Funding acquisition, Balpreet Ahluwalia and Peter McCourt; Investigation, Hong Mao, Karolina Szafranska and Glen Lock-wood; Methodology, Hong Mao, Karolina Szafranska, Cynthia Whitchurch, Robin Diekmann, David Le Couteur and Victoria Cogger; Project administration, Peter McCourt; Resources, Cynthia Whitchurch, Louise Cole and Peter McCourt; Software, Deanna Wolfson and Robin Diekmann; Supervision, Peter McCourt; Validation, Hong Mao and Karolina Szafranska; Visualization, Hong Mao, Karolina Szafranska and Deanna Wolfson; Writing – original draft, Hong Mao and Karolina Szafranska; Writing – review & editing, Hong Mao, Karolina Szafranska, Larissa Kruse, Christopher Holte, Deanna Wolfson, Balpreet Ahluwalia, Cynthia Whitchurch, Louise Cole, Glen Lockwood, Robin Diekmann, David Le Couteur, Victoria Cogger and Peter McCourt. All authors have read and agreed to the published version of the manuscript.

## Acknowledgments

The authors would like to thank Randi Olsen and Tom-Ivar Eilertsen from Advanced Microscopy Core Facility at UiT for the electron microscopy expertise.

