## Supplemental information for "Effect of caffeine and other xanthines on liver sinusoidal endothelial cell ultrastructure"

**Supplementary information**

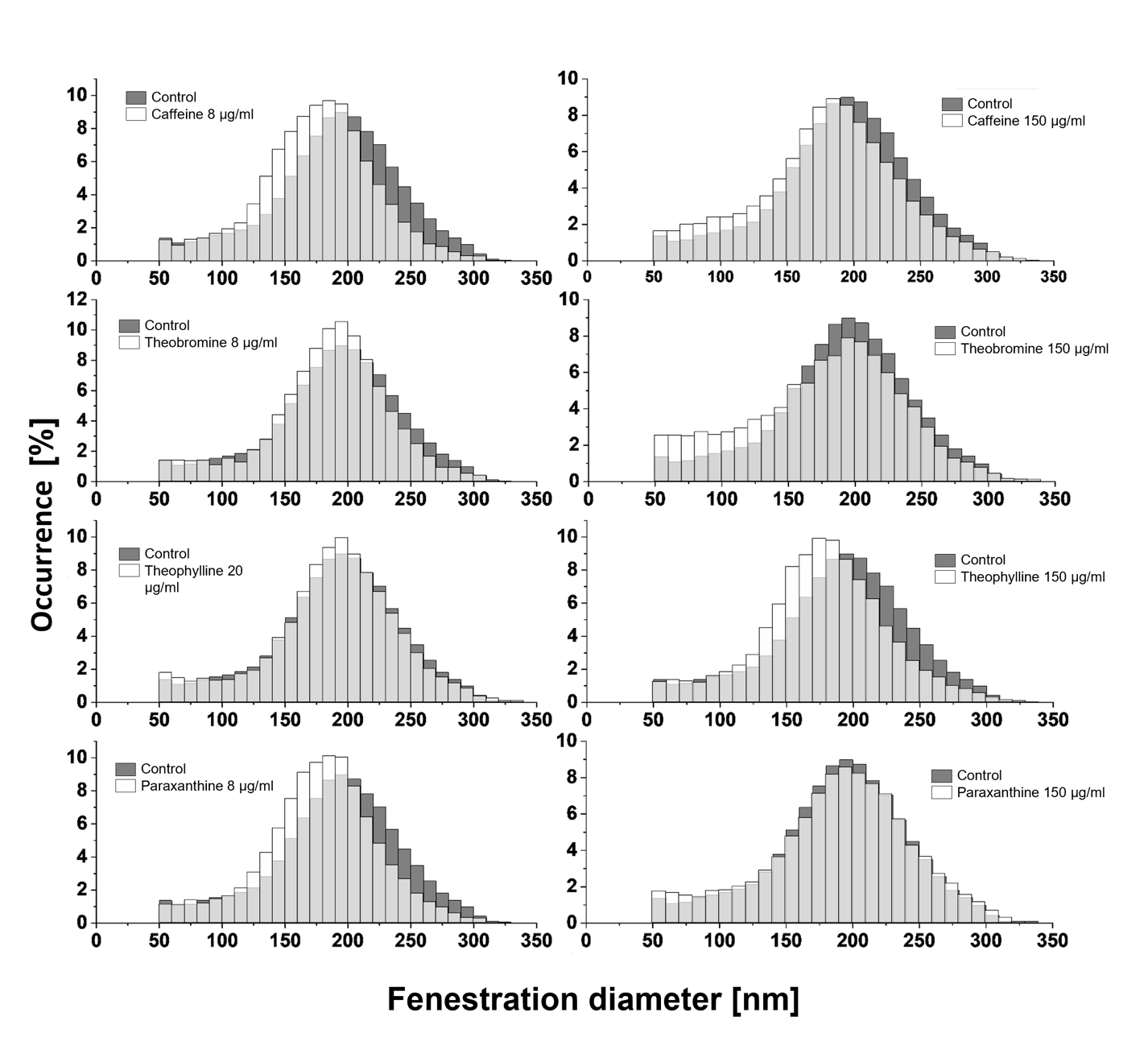
S1. Fenestration size distributions

**Figure S1.** Distribution of fenestration size after treatment, in comparison with control/untreated samples

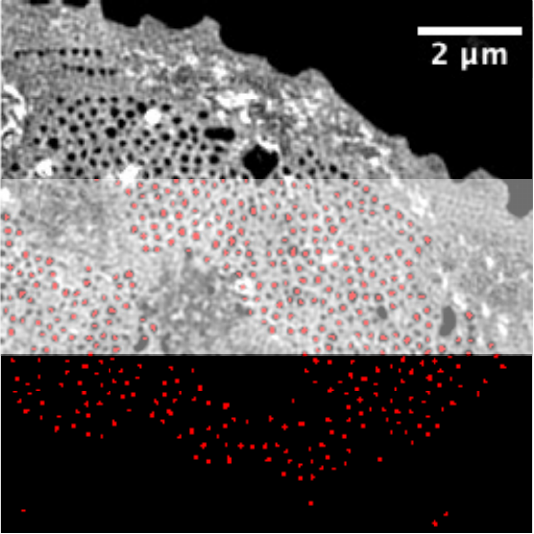

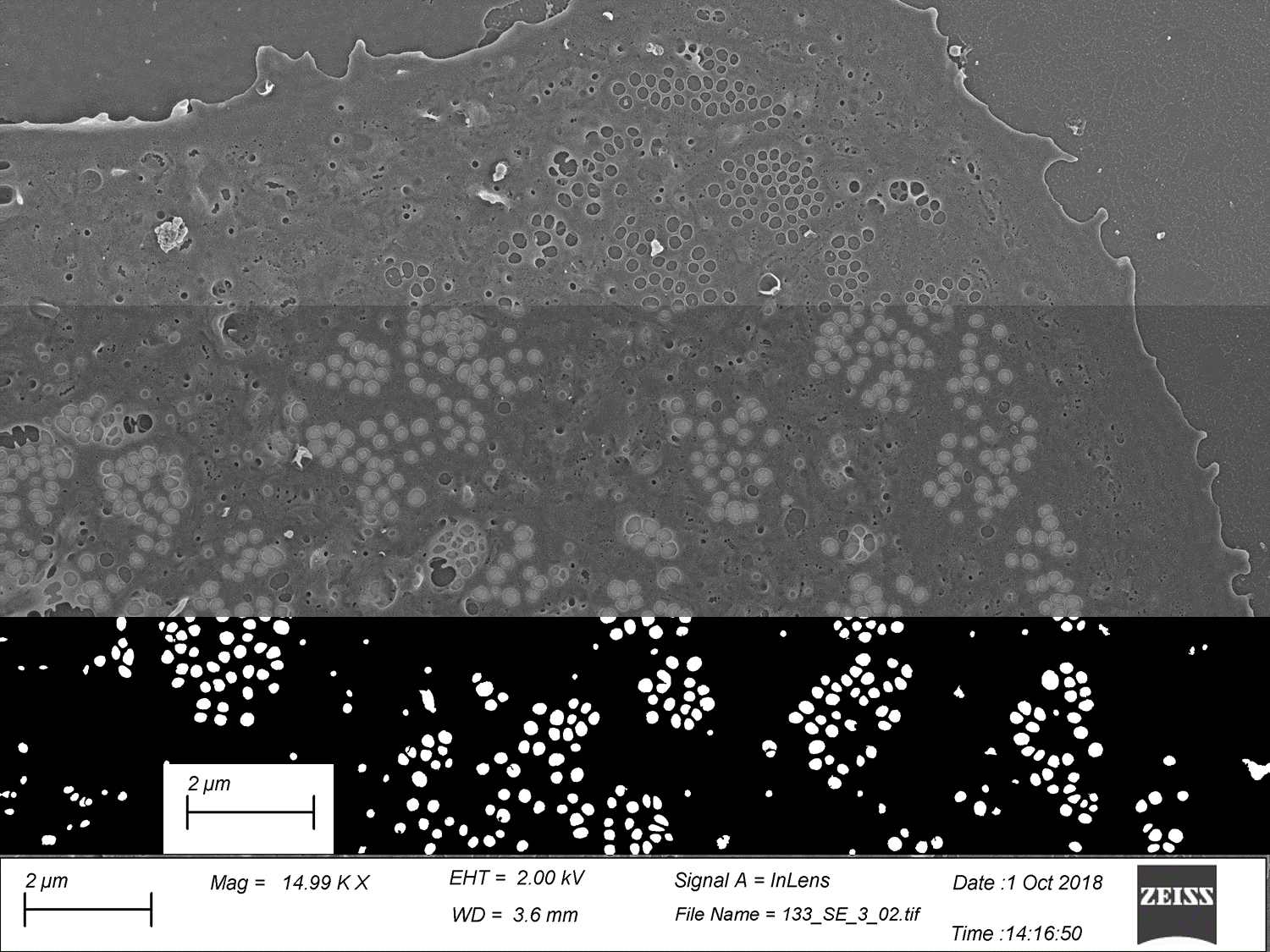
S2. Image analysis data

**Figure S2.** Examples of the images used for measurement of fenestration diameters. The top section is a raw SEM (left) / SIM (right) image, bottom section presents the binary mask used for quantitative analysis, and the middle section shows the overlay used to identify fenestrations.
